# Sensitivity profiling reveals consistent drug responses across preclinical neuroblastoma models

**DOI:** 10.64898/2026.02.24.707695

**Authors:** Marlinde. C. Schoonbeek, Marloes van Luik, Heike Peterziel, Dennis Gürgen, Vicky Amo-Addae, Lindy Vernooij, Eleonora J. Looze, Sabina Valova, Sina Kreth, Jan Koster, Angelika Eggert, Julia Schueler, Aniello Federico, Apurva Gopisetty, Benjamin Schwalm, Marcel Kool, Alexandra Saint-Charles, Gudrun Schleiermacher, Gilles Vassal, Hubert N. Caron, Lou Stancato, Stefan M. Pfister, Frank Westermann, Jens Hoffman, Arjan Boltjes, Marlinde L. van den Boogaard, Selma Eising, Ina Oehme, Jan J. Molenaar, Sander R. van Hooff

**Author notes:** Correspondence in submission process should be addressed to M. C. Schoonbeek, Princess Máxima Center, Utrecht, the Netherlands. Correspondence after publication should be addressed to J.J. Molenaar (; lead contact), Princess Máxima Center, Utrecht, the Netherlands.

## Abstract

Despite intensive treatment, overall survival for high-risk and relapse neuroblastoma patients remains below 50%. Even though comprehensive molecular profiling enables treatment stratification, druggable alterations have been identified for only a subset of patients. *In vitro* drug screening offers a complementary approach. Here, we compare the translational potential of three preclinical drug screening methods: *ex vivo* short-term, *in vitro* patient-derived organoid and *in vivo* patient-derived xenograft (PDX) drug testing. In total, 55 screens were performed from 38 neuroblastoma samples and five pediatric non-malignant samples, testing 77-224 drugs per screen. *Ex vivo* short-term drug screens achieved higher success rates than organoid screens (65% versus 23%) and shorter turnaround times (14 days versus 3-12 months). Matched samples showed consistent drug sensitivities across sample origin (patient versus PDX-derived; mean r = 0.84) and method (*ex vivo* short-term versus organoids; mean r = 0.87), demonstrating that *ex vivo* short-term screens recapitulate drug sensitivities found in long-term organoid models. In parallel, as part of the ITCC-P4 consortium, ten compounds were tested *in vivo* in eight PDX models, with samples matching the *ex vivo* screens. For seven out of ten clinically available compounds, *ex vivo* drug responses were comparable with *in vivo* responses in matched PDX models. These results demonstrate that, while organoids and PDX models remain essential for drug discovery, *ex vivo* short-term drug screening provides a rapid alternative for functional precision oncology in neuroblastoma.

## Introduction

Neuroblastoma arises from the developing sympathetic nervous system and remains one of the most challenging pediatric solid tumors to treat. Despite advances in treatment, effective therapy options remain limited for high-risk and relapsed neuroblastoma, with overall survival rates at ∼50%.^1–3^ Pediatric precision oncology programs have been established with the aim of identifying druggable tumor driving alterations through molecular profiling.^4–6^ Although promising, for many patients no targetable alterations could be identified, and the clinical translation was limited.^4–6^ This highlights the need for additional strategies in personalized medicine to guide treatment selection.

As a complementary approach to molecular profiling, drug screening can be used to guide treatment.^7^ For this purpose, *in vivo* Patient-Derived Xenografts (PDXs) and *in vitro* organoid models have been used in neuroblastoma.^8–10^ However, these methods are time-consuming, and expensive to develop and maintain.^9,11^ Moreover, the clinical application of these models is further impeded by low establishment rates and long establishment times, which often exceed the median time to progression of four months, as we reported previously.^12^ *Ex vivo* short-term drug screening (in short: short-term screens), where freshly obtained tumor tissue is briefly cultured before drug screening, offers a rapid alternative, delivering drug response profiles within a clinically relevant timeframe of 7-24 days after sample collection.^13–15^

In this study, we evaluated integrated drug sensitivity and molecular profiling in different neuroblastoma models to assess their consistency in drug sensitivity and their potential to guide personalized treatment selection. We performed short-term and organoid screens and combined this with our previous work on patient-derived organoids,^9^ *ex vivo* short-term screens derived from patients^14^ or PDX models.^15^ In parallel, *in vivo* compound testing on matched PDX models was performed within the ITCC-P4 initiative, a European public-private partnership evaluating preclinical drug efficacy in pediatric cancer PDX models.^16,17^ Furthermore, we included pediatric non-malignant samples to assess non-selective drug toxicity. Our results demonstrate consistent drug sensitivities across different neuroblastoma models, assay durations and tissue sources, and highlight the translational potential of *ex vivo* short-term drug screening.

## Results

### Establishment and characterization of short-term, organoid, and PDX neuroblastoma models

To compare drug sensitivities of pre-clinical neuroblastoma models, 55 drug screens, from samples originating from 38 patients, were performed at the Princess Máxima Center (Máxima) or Hopp Children’s Cancer Center (KiTZ) (Fig. 1A; Suppl. Fig. 1A). Four types of drug screens were performed on tumor cells: patient-derived short-term (n = 8) and organoid (n = 13) screens, and PDX-derived short-term (n = 17) and organoid (n = 8) screens (Fig. 1B; Suppl. Fig. 1B). In addition, five short-term screens were performed on pediatric non-malignant tissue, including fibroblasts (n = 2), astrocytes (n = 2), and PBMCs (n = 1). The feasibility of the different approaches was assessed by comparing establishment timelines and success rates (defined as successful culture establishment and high-quality drug screen). Short-term screens of both patient and PDX material provided drug sensitivity profiles within 14 days of sample receipt, whereas organoid models required 3-12 months. In the Máxima cohort of PDX-derived samples, short-term screens were successful in 13/20 cases (65%), whereas organoid screens succeeded in 5/22 cases (23%). Failures in short-term screens were mainly due to insufficient viable cell yield (n = 6) or screen quality (n = 1), while failures in organoid screens were all due to growth arrest during long-term culture (n = 17). Of note, four screens were excluded from further analyses due to murine content (n = 2) or because they represented duplicate screens from the same patient (n = 2).^15^ These results show that both the turnaround times and success rates of short-term screens are superior to organoid screens, indicating that short-term screens could be more suitable for real-time clinical decision making.

**Figure 1.**
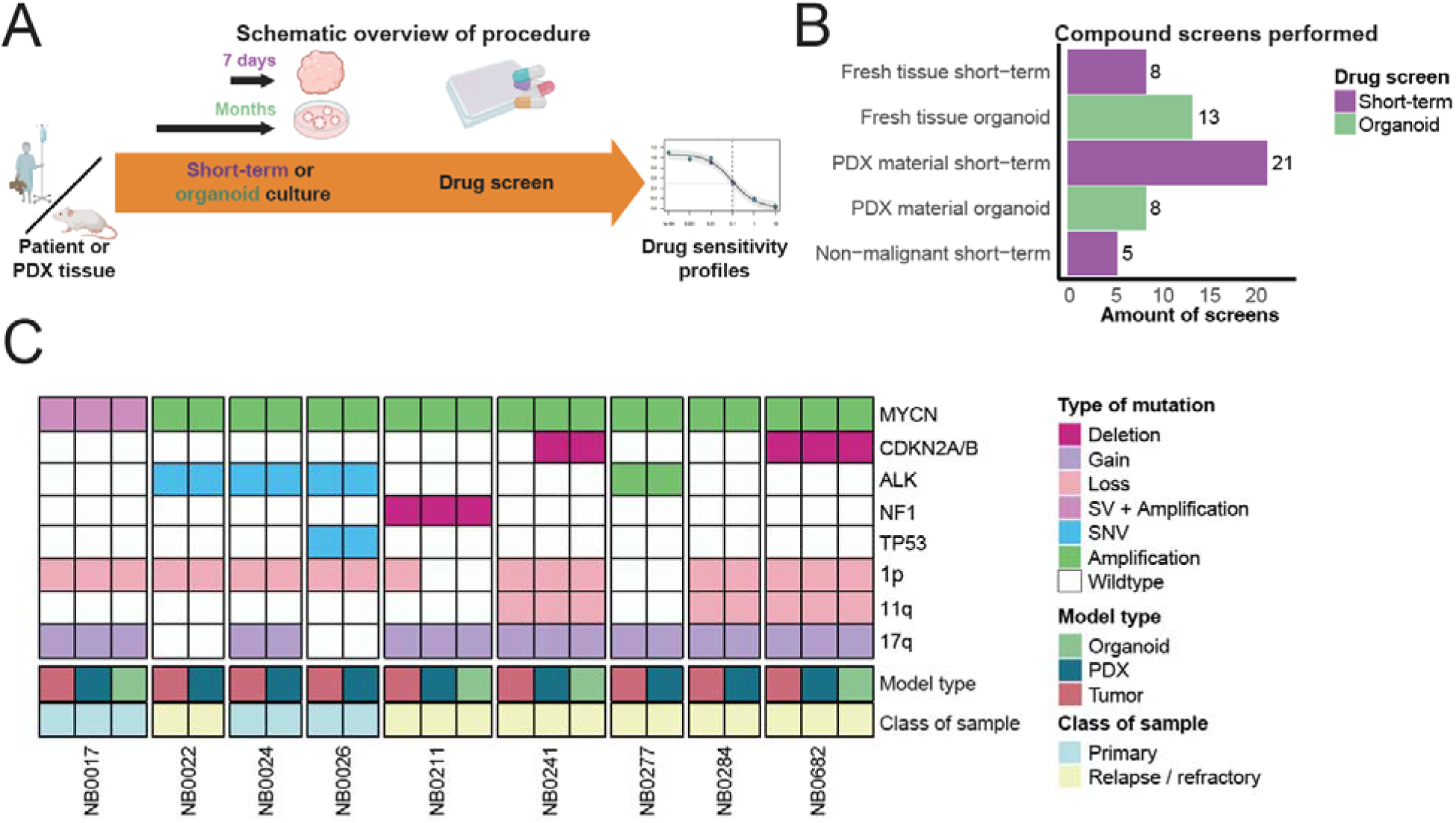
Establishment and genetic characterization of models for drug screening. **A)** A schematic overview of model establishment and drug screening. Samples were obtained from patients or PDX models, digested, and cultured either for seven days (short-term) or until stable growth was observed (months; organoids) before dru screening. Drug response was determined using the area under the curve. **B)** Bar plot representing the total number of drug screens per model type (purple: short-term; green: organoid). Part of the short-term and organoid screens performed at KiTZ were previously described in Peterziel et al.^14^, Máxima patient material organoids were previously described in Langenberg et al.^9^, and short-term screens performed at Máxima in Schoonbeek et al.^15^. **C)** Oncoplot of genomic landscape of matched neuroblastoma samples from Máxima, showing key gene alterations and chromosomal aberrations colored by alteration type. ALK mutations include: F1174L (NB0022) and R1275Q (NB0024 and NB0026). PDX = Patient-Derived Xenograft; SNV = single-nucleotide variant; SV = structural variant.

To interpret drug sensitivity profiles in a molecular context, we analyzed whole exome sequencing, RNA sequencing and methylation array on patient tumors, PDX tumors, and the organoid models for which drug screens were performed in the Máxima. The most frequent molecular events were *MYCN* amplification, *ALK* mutation, loss of chromosome 1p, loss of chromosome 11q, and gain of chromosome 17q (Suppl. Fig. 1C). To determine if PDX and organoid models retain the genetic alterations of the originating patient tumor, we examined driver events in models where tumor data was available. In matched samples, all driver events were maintained with the exception of a chromosome 1p loss observed in the NB0211 patient tumor but absent in the PDX and organoid models (Fig. 1C). We also assessed whether gene expression patterns were consistent for organoids in the Máxima cohort. We indeed show that models generally cluster close to the originating tumor (Suppl. Fig. 2). These results demonstrate that neuroblastoma short-term, organoid, and PDX models can be successfully established while maintaining the key genomic and phenotypic features of the original patient tumor.

### Consistent drug sensitivities across patient- and PDX-derived models, and across short-term and organoid models

To evaluate drug sensitivities across preclinical neuroblastoma models, screening was performed with drug libraries of 77-224 pediatric cancer-approved and clinical trial compounds, of which 60 were shared between Máxima and KiTZ (Fig. 2A; Suppl. Table 3). The drug libraries included cytotoxic and targeted therapies, tested at concentrations ranging from 1 x 10^-6^ μM to 250 μM (Suppl. Fig. 3). Drug response was quantified using the area under the curve (AUC), ranging from 0 (sensitive) to 100 (resistant) (Suppl. Fig. 4-5).

**Figure 2.**
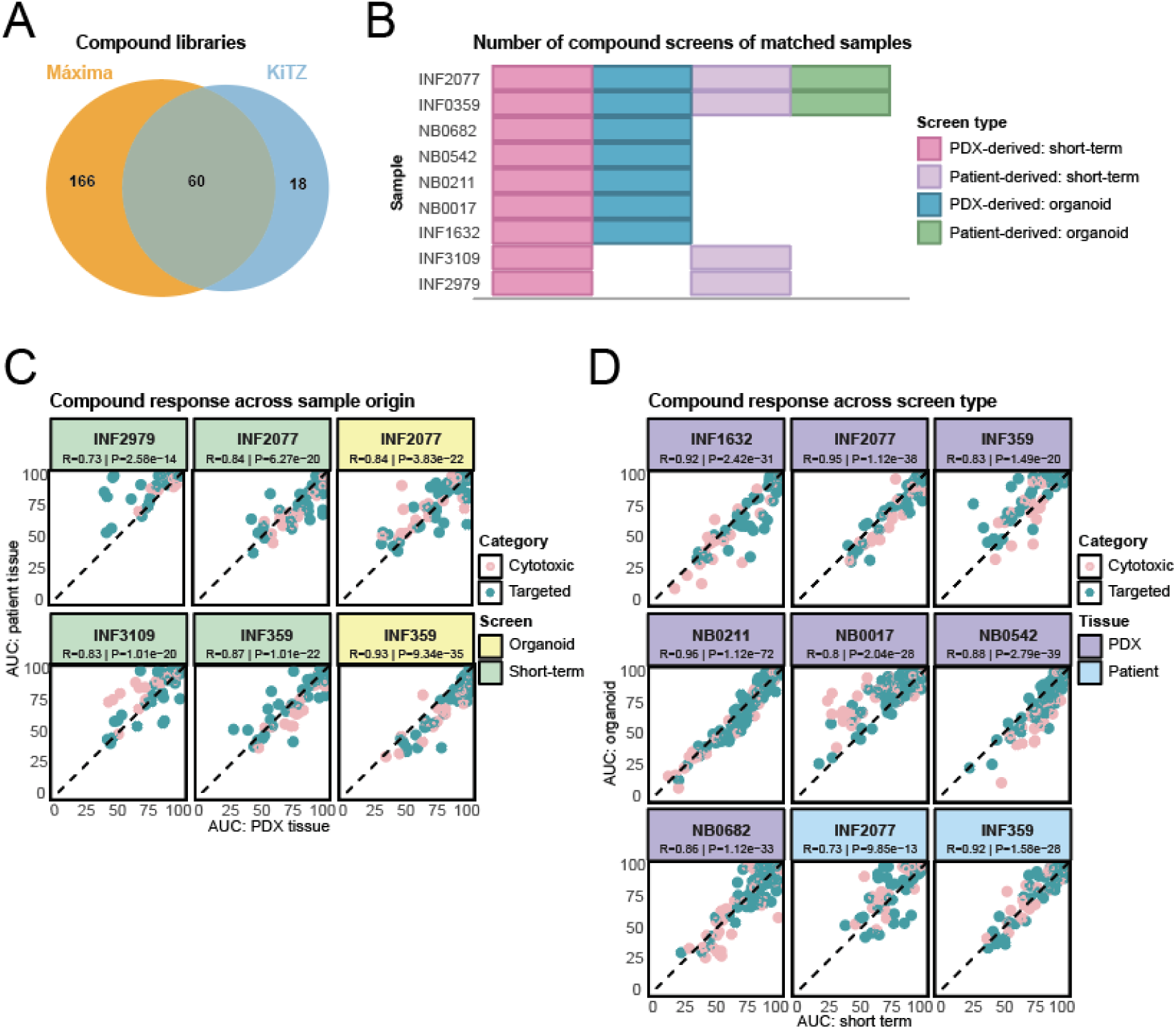
Comparison of drug response across neuroblastoma model systems and sample origin. **A)** Overlappin drugs of the Máxima and KiTZ libraries. **B)** Screens performed on matched samples, colored by their model. **C)** Correlation plots of drug response (AUC) on organoid or short-term screens from matched samples derived from different tissue origins (patient or PDX material). Each dot represents a drug, colored by category: cytotoxic or targeted. A lower AUC defines higher sensitivity. **D)** Correlation plots of the drug response (AUC) of organoids an short-term cultures established from matched samples. Each dot represents a drug, colored by category: cytotoxic or targeted. A lower AUC defines higher sensitivity. AUC = area under the curve; R = Pearson correlation; P = p-value.

To assess whether drug sensitivity is influenced by *in vivo* expansion, we compared drug response of six matched samples, including short-term (n = 4) and organoid (n = 2) screens from either patient or PDX material (Fig. 2B). Matched drug response profiles are highly correlated despite differences in sample origin (mean r = 0.84; Fig. 2C; Suppl. Fig. 6; Suppl. Fig. 7A). No drug was significantly associated with origin of the model indicating that this factor did not have a major influence on drug sensitivity (Suppl. Fig. 7B). These results demonstrate that PDX-derived models recapitulate drug sensitivity observed in patient-derived material.

To evaluate the influence of the duration of *in vitro* culturing (max. 7 days versus 3-12 months), we compared drug sensitivities of matched short-term and organoid screens (PDX n = 7, patient n = 2). In the Máxima cohort, additional methodological differences between short-term and organoid screens existed: the culture medium and the compound incubation time (72 hours versus 120 hours in the organoids). Despite these differences, short-term and organoid screens show similar trends in drug response (average Pearson correlation = 0.87; Fig. 2D; Suppl. Fig. 7C), and no drugs significantly differed in their efficacy across the two model systems (Suppl. Fig. 7D-E). Next to consistent drug sensitivities across patient- and PDX-derived tissues, these results indicate that short-term screens recapitulate drug responses of the organoids, the current standard for functional profiling,^9,18,19^ while shortening the turnaround from months to days.

### Genomic and transcriptomic features in relation to *in vitro* drug response

To evaluate whether genomic or transcriptomic features of untreated neuroblastoma samples are associated with drug sensitivity in short-term and organoid models, we associated drug response with corresponding molecular data. When multiple screens of one sample were available, one representative screen (the short-term PDX-derived screen, available for all these samples) was included in this analysis. To evaluate whether targetable genetic alterations are associated with *ex vivo* drug response, we compared sensitivities in altered versus wild-type samples. Across the two cohorts, three targetable alterations were significantly associated with drug response (Fig. 3A; Suppl. Fig. 8A). Increased sensitivity to MDM2 inhibition was observed in TP53 wild-type samples compared to homozygous TP53-altered samples, with a significant association for idasanutlin and XPO1 inhibitor selinexor in the Máxima cohort, and trends in the same direction in the KiTZ cohort. These findings validate previous reports of resistance of tumors with a TP53 inactivation to MDM2 and XPO1 inhibitors.^20,21^ Next to this, ALK alterations are significantly associated with sensitivity to ALK inhibitor lorlatinib in the Máxima cohort, with the same trend in KiTZ. However, ALK alterations were not associated with sensitivity to other ALK inhibitors, and other alterations such as SMARCA4 and MYCN were not significantly associated with drug sensitivity in either cohort. These results show that while some genetic alterations, such as a major tumor driver like TP53, associate with *ex vivo* drug response, many genetic alterations alone may not be enough to predict *ex vivo* drug sensitivity in neuroblastoma.

**Figure 3.**
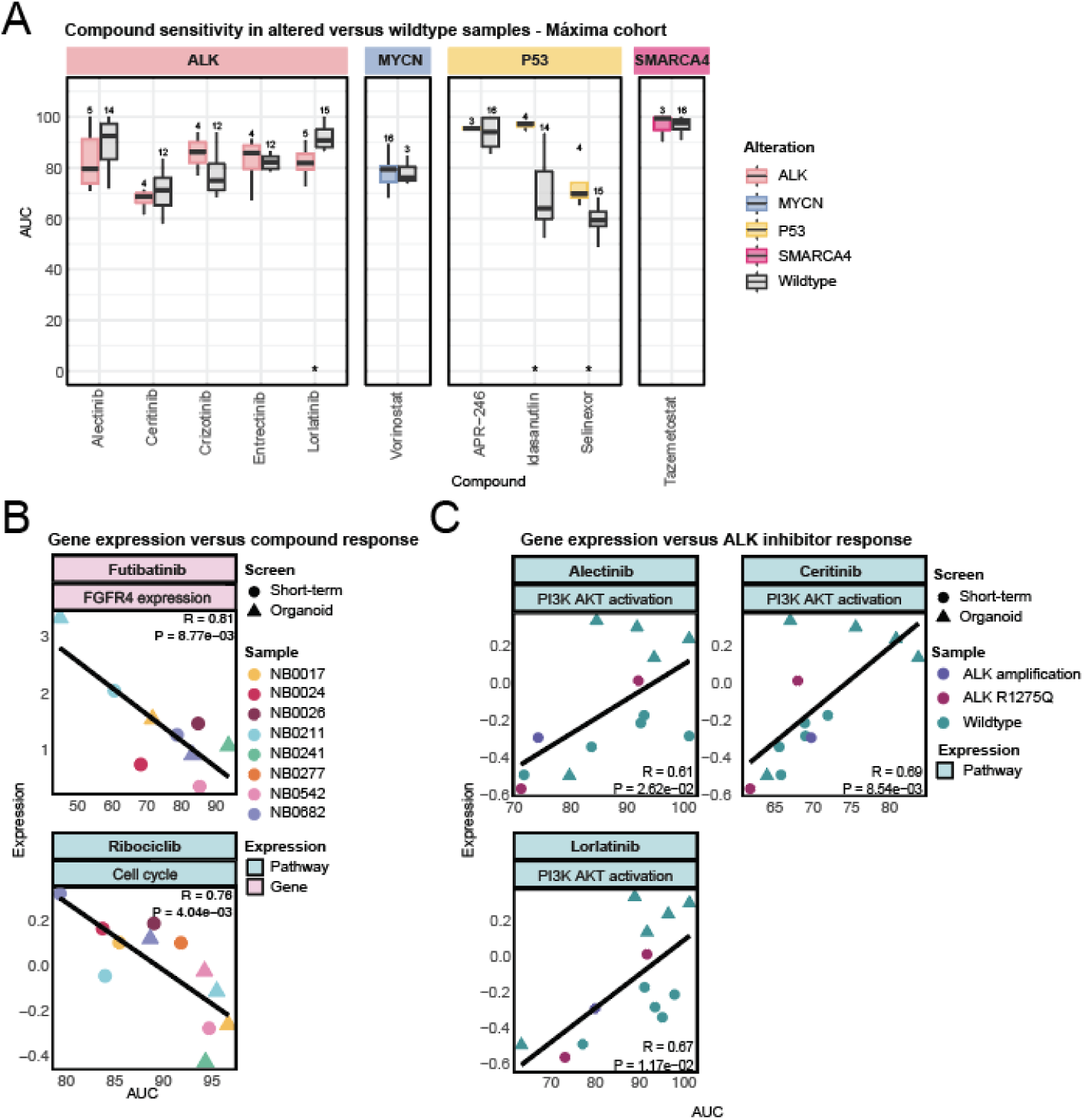
Genomic and transcriptomic features associations with in vitro drug sensitivity. **A)** AUC values of samples with targetable gene alterations, in boxes colored by alteration, compared to wildtype samples, in grey boxes, of the Máxima cohort. Statistical testing was only possible when there were at least three samples in group, and stars indicate a significant difference between the altered and wildtype samples (adjusted p-value < 0.05). **B)** Correlation of AUC of futibatinib with FGFR4 expression and ribociclib with the cell cycle pathway, each dot represents a sample, colored by name and shaped by the type of screen (short-term or organoid). C) Correlation of AUC of ALK inhibitors alectinib, ceritinib, and lorlatinib with the enrichment of the targeted pathway PI3K AKT. Eac dot represents a sample, colored by alteration. AUC = area under the curve; R = Pearson’s correlation.

To explore whether expression of target genes or pathways reflects drug sensitivity, drug responses were correlated with RNA expression of each compound’s known target gene or pathway, where transcriptomic data were available (3B-C; Suppl. Fig. 8B). The strongest correlations included an association between sensitivity to futibatinib and higher FGFR4 expression and sensitivity to ribociclib with elevated expression of cell cycle genes (Fig. 3B).

For only one out of five ALK inhibitors, we observed an association between ALK status and sensitivity to ALK inhibition, despite the use of ALK status as a biomarker for ALK inhibition in clinical trials.^22^ Therefore, we additionally looked into expression of ALK and the PI3K pathway. We did not find a significant correlation between ALK expression and ALK inhibitor sensitivity, apart from entrectinib (Suppl. Fig. 8B), driven by one ALK-amplified sample with elevated ALK expression. However, reduced activity of the PI3K-AKT pathway did correlate with increased sensitivity to ceritinib, alectinib, and lorlatinib (Fig. 3C). The PI3K-AKT pathway functions downstream of ALK signaling and has been previously reported as a resistance mechanism.^23,24^ The observed associations highlight that RNA expression can provide additional insights into drug sensitivities beyond mutational status.

### Drug sensitivity profiling to assess tumor selectivity in neuroblastoma

To identify if drugs are more effective in tumor or non-malignant cells, we compared the drug sensitivities of neuroblastoma to non-malignant pediatric samples (fibroblasts (n = 2), astrocytes (n = 2), and PBMCs (n = 1)) screened at KiTZ. First, we analyzed the compounds comprising the clinical backbone of high-risk neuroblastoma treatment as used in the SIOPEN HR-NBL2 trial^25^ (Fig. 4A). Cisplatin (p = 0.04) and etoposide (p = 0.19) were more effective in tumor than non-malignant samples. In contrast, doxorubicin and vincristine were similarly effective.

**Figure 4.**
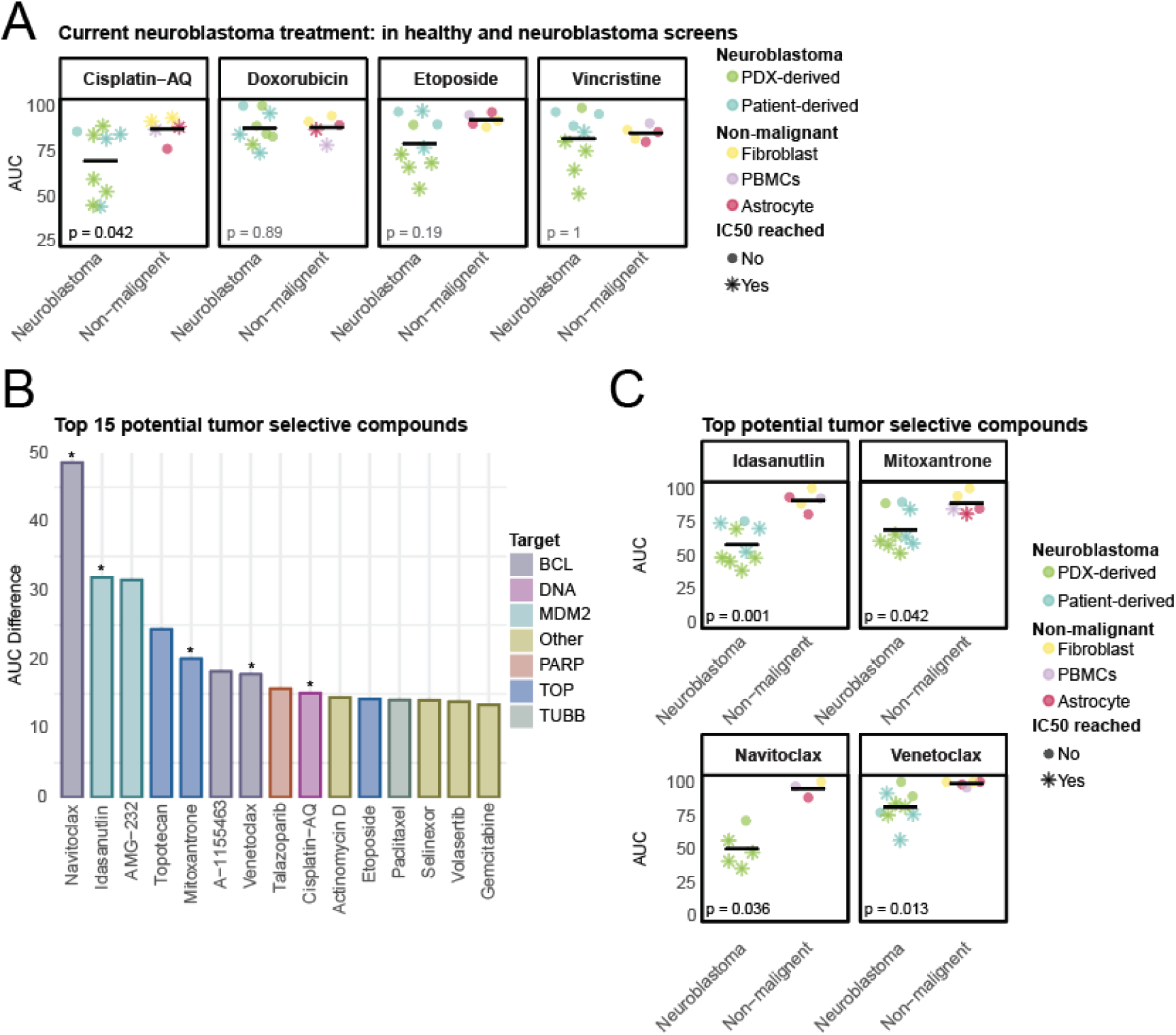
Drug screening of neuroblastoma and pediatric non-malignant samples in the KiTZ cohort. **A)** Box plots representing drug sensitivity (AUC) of tumor and non-malignant samples for compounds currently used in the clinic. Each dot represents the drug response of one sample, colored by tissue type. Stars indicate that the IC50 was reached within the tested concentrations. The line indicates the mean AUC. **B)** Bar plot representing the top 15 tumor selective drugs. The top 15 drugs with the highest mean AUC difference between tumor and non-malignant samples (mean AUC non-malignant – mean AUC tumor) are shown. Bars are colored based on the target of th drug. Asterisks indicate that the drug is significantly more effective in tumors. **C)** Plots representing drug sensitivity (AUC) of tumor and non-malignant samples for drugs that were significantly more effective in neuroblastoma. Each dot represents the drug response of a sample, colored by tissue type. Stars indicate that the IC50 was reached within the tested concentrations. The horizontal line indicates the mean AUC of the samples. AUC = area under the curve.

To identify tumor-specific drugs, we compared drug sensitivity in tumor and non-malignant models. Figure 4B shows the 15 most tumor selective drugs. Besides cisplatin (Fig. 4A), four compounds were significantly more effective in tumor samples: BCL2 inhibitors navitoclax and venetoclax, MDM2 inhibitor idasanutlin, and topoisomerase inhibitor mitoxantrone (Fig. 4C). Conversely, we identified drugs for which non-malignant cells are more sensitive (Suppl. Fig. 9A): MEK inhibitors cobimetinib and trametinib, tyrosine kinase inhibitor dasatinib, mTOR inhibitors everolimus and temsirolimus, and HDAC inhibitor panobinostat (Suppl. Fig. 9B). These findings highlight the value of including pediatric non-malignant tissues as a reference in drug screens to identify compounds with favorable tumor selectivity and flag those with potential toxicity.

### Concordance between *ex vivo* and *in vivo* drug responses in preclinical neuroblastoma models

*In vivo* compound studies were performed within the ITCC-P4 consortium.^16,17^ For every compound a number of different PDX models were tested in a single mouse trial setup to optimize the number of tested compounds and models while limiting animal numbers. Ten compounds were tested in eight neuroblastoma PDX models (Fig. 5A; Suppl. Fig. 10A; Suppl. Table 4). *In vivo* response was determined as end-of-treatment relative tumor volume, normalized to the starting tumor volume.

**Figure 5.**
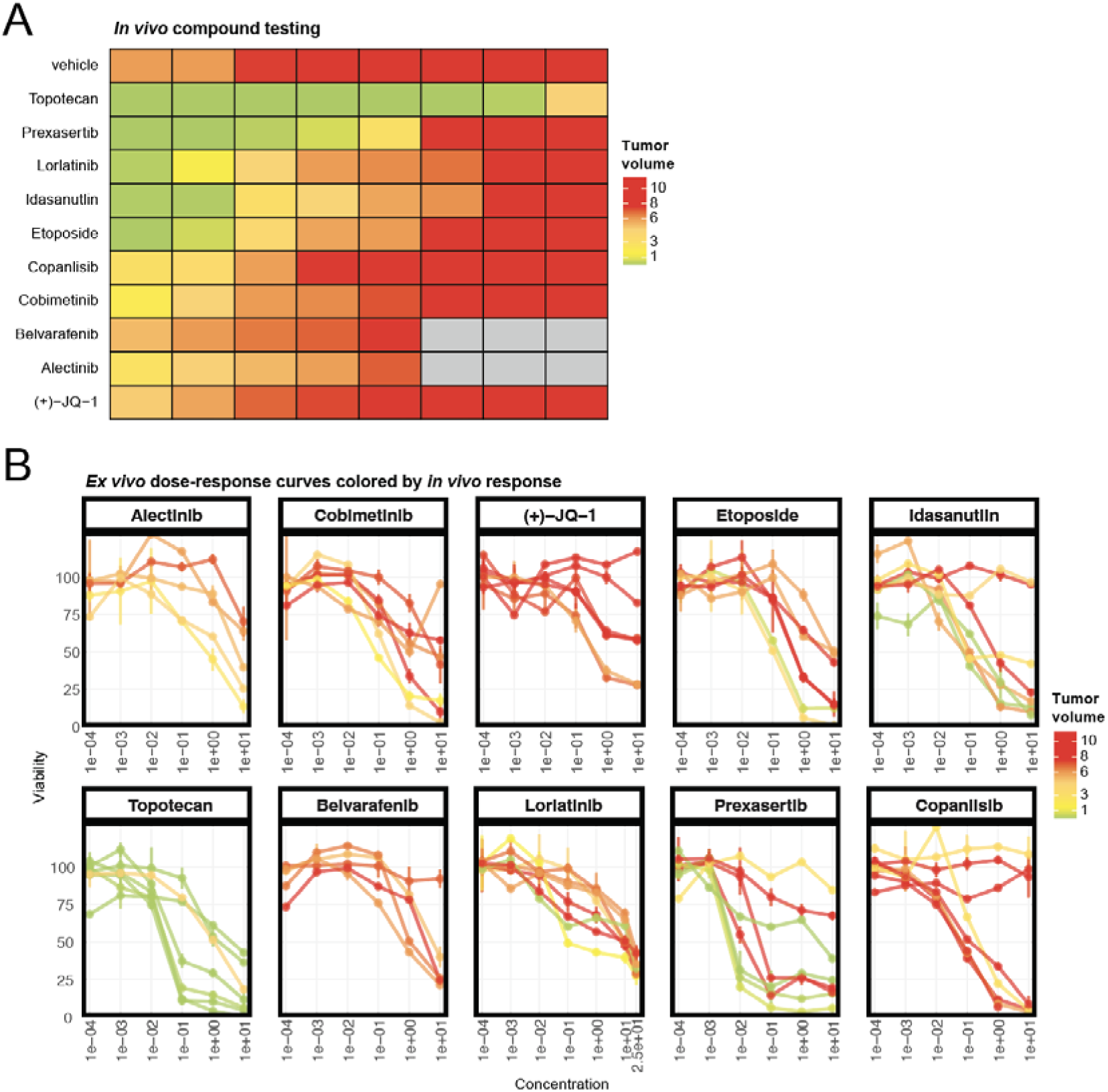
PDX treatment response reflects ex vivo response. **A)** Heatmap representing the in vivo treatment responses of PDX models. Each row represents a compound (ordered by response), colors indicate end-of-treatment response (tumor volume, normalized to starting volume). **B)** Ex vivo dose response curves. Each line represents the ex vivo dose response curve of a sample and is colored by in vivo end-of-treatment-response (tumor volume) in the matched PDX model. Each dot represents the viability of the cells at each concentration, with an error bar showing the variability of the two replicates (if available). Compounds are ordered by spearman correlation (AUC versus tumor volume; left top: highest spearman rho = 1; bottom right: lowest spearman rho = 0.05).

To evaluate concordance between drug sensitivity in *ex vivo* short-term screens and therapeutic responses in corresponding PDX models, we compared the *ex vivo* AUCs with *in vivo* tumor volume. Although the assays differ substantially in design (constant 72-hour exposure across six concentrations *ex vivo* versus intermittent dosing of a single concentration over 28 days *in vivo*), we observed concordant response patterns for several compounds. Topotecan was highly effective in both model systems, reducing *in vivo* tumor volume and *ex vivo* viability in most samples (Suppl. Fig. 10A-B). In contrast, belvarafenib was generally not effective in both settings, with high tumor volumes *in vivo* (tumor volumes >5) and low *ex vivo* sensitivity (AUC > 75). As compounds that are effective for only part of the patients are most relevant for precision oncology, we assessed sample-level concordance (Fig. 5B; Suppl. Fig. 10C). Three drugs showed strong rank-based concordance (spearman ρ > 0.6): the ALK inhibitor alectinib (ρ = 1), the MEK inhibitor cobimetinib (ρ = 0.71) and the chromatin-remodeling BET inhibitor JQ-1 (ρ = 0.61). Moderate concordance was observed for the topoisomerase inhibitor etoposide (ρ = 0.54) and the MDM2 inhibitor idasanutlin (ρ = 0.45). For idasanutlin, most *ex vivo* responders also responded *in vivo.* However, an exception was NB0542, which was resistant *in vivo* (tumor volume = 17), despite moderate ex vivo sensitivity (AUC = 76, slightly above the mean of 72) – consistent with its TP53 wild-type status. Weak concordance (ρ < 0.4) is found for three compounds: copanlisib, prexasertib and lorlatinib. Overall, ex vivo sensitivity patterns reflected *in vivo* PDX response for seven out of ten drugs tested.

## Discussion

Despite intensive treatment, survival of high-risk neuroblastoma patients remains around 50%. Molecular profiling alone is insufficient to stratify treatment for all patients.^12,26–28^ Functional profiling using organoids and patient-derived xenograft (PDX) models has shown promise but is limited by establishment timelines and success rates.^9,11,12^ Here, we evaluated the suitability for precision medicine of three types of patient-derived neuroblastoma models: the short-term cultures, organoids, and PDX models. We performed 55 drug screens on tissues derived from 38 neuroblastoma patients and 5 non-malignant samples. In parallel ten compounds were tested *in vivo*, in eight neuroblastoma PDX models.^17^ Short-term drug screens achieved higher success rates and shortened establishment times compared to organoids while retaining the drug response profiles. Short-term screens also showed concordance with *in vivo* PDX model treatment response for a subset of compounds.

PDX models remain widely used in neuroblastoma to enable *in vivo* assessment of drug efficacy and tolerability.^8,9^ In our study, PDX models retained the major genetic and transcriptomic characteristics of their originating tumors. However, their suitability for personalized precision medicine is constrained by low engraftment success, high costs and long establishment times.^29,30^ Moreover, treatment responses in PDXs do not always fully mirror patient activity or toxicity, as murine pharmacokinetics and pharmacodynamics differ from those in children.^29,30^ In addition, the use of animals raises ethical considerations, and regulatory agencies such as the FDA increasingly encourage alternatives to animal-based preclinical testing.^31^ These limitations motivate the search for alternative patient-derived model systems that are eligible for precision medicine.

We compared four different *ex vivo* drug screening approaches and observed concordance in the responses of short-term and organoid screens, as well as between drug screens derived from patient material and those derived from PDX. To the best of our knowledge, such cross-model comparisons have rarely been systematically explored. Previous work from the ZERO program also demonstrated strong concordance between short term screens of matched patient- and PDX-derived tissue for a single rhabdomyosarcoma case.^13^ Our study extends on these observations and provides a unique dataset which allows direct comparison of drug responses before and after *in vitro* and/or *in vivo* expansion.

Organoid models can be derived from both patient biopsies and PDXs and are attractive for large-scale drug discovery and mechanistic studies. In our cohort, PDX-derived neuroblastoma organoids required 3-12 months to establish with a 23% success rate. Success rates for patient biopsies are likely lower due to limited biomass. Similar timelines have been reported by others for neuroblastoma organoid models,^9,10,32^ often exceeding the median progression-free survival of relapsed or high-risk patients (4 – 7 months), which restricts their use for real-time clinical decision making.^11^ A second potential downside is clonal selection during prolonged culture. Karlsson *et al*.^32^ demonstrated that within 42 days (11 passages), the major subclone can expand to fixation while others are lost. Such evolution over months of culture may influence drug responses by not fully reflecting the polyclonal composition of the tumor. Overall, these factors limit the suitability of organoids for real-time clinical decision making.

*Ex vivo* short-term cultures offer a faster approach for preclinical drug testing. In our study, short-term screens derived from PDX models achieved a 63% success rate and generated drug-response profiles within 14 days. This aligns with our previously reported patient-derived success rate (58%^14^). *Ex vivo* short-term screens can be used for precision medicine, by comparing individual drug responses to the cohort to identify patient-specific outlier sensitivities.^15^ Recently, ZERO and INFORM demonstrated the feasibility of integrating *ex vivo* short-term screens cultures in precision medicine programs as the rapid turnaround time allows integration into multidisciplinary tumor boards.^13,14^ In ZERO, 82% of screens were completed while patients were still alive. In addition, in INFORM, we observed preservation of the tumor microenvironment (with a trend for tumor cell enrichment) during preculture and drug screening. As short-term cultures can retain residual microenvironmental cells, they may provide a more representative tumor model than long-term models. To account for residual microenvironment cells, future studies should develop methods to distinguish between tumor and non-tumor cells to determine the tumor-specific drug response. In addition, further optimizing success rates and reducing the required cell numbers per well will be essential for implementation of drug screening when using small needle biopsies.

To assess whether *ex vivo* drug responses correspond with *in vivo* activity, we compared *ex vivo* drug sensitivity from short-term screens with matched PDX model responses, performed within the ITCC-P4 consortium.^16,17^ We observed moderate to strong concordance for seven agents, including kinase inhibitors targeting ALK (alectinib), MEK (cobimetinib) and MDM2 inhibitor idasanutlin. For others (e.g. copanlisib), concordance was limited, which could be due to several factors. Direct comparisons are constrained by major differences between the models, such as discordant drug concentrations, treatment durations, and pharmacokinetic and pharmacodynamic parameters.^33^ For rapidly metabolized compounds, *ex vivo* concentrations may be higher than concentrations achievable in plasma or tumor tissue, which may explain the lack of *in vivo* efficacy even when *ex vivo* sensitivity is observed. Another potential difference is the discrepancy between *in vitro* and *in vivo* growth rates, which especially impacts sensitivity to cytotoxic agents^34,35^. Besides these differences, *in vivo* models are known for their biological variability in treatment response.^36^ Therefore, using an n-of-1 setup limits our direct comparison between *in vivo* and *ex vivo* sensitivity. Lastly, the PDXs tested *in vivo* were not from the same passage from which the *ex vivo* models were derived, introducing the possibility of clonal shifts between the passages. Ultimately though, direct correlation with in-patient, not PDX, response holds clinical value. Previous work by ZERO^13^ and INFORM^14^ demonstrated that *ex vivo* sensitivities were consistent with responses in matched PDXs and, critically, in patients.

Beyond predicting tumor response, functional assays may also provide insights into potential tumor selectivity and toxicity. Two of the clinical backbone agents used in high-risk neuroblastoma treatment, doxorubicin and vincristine, showed comparable effectiveness in neuroblastoma and non-malignant samples. This is consistent with previous reports of doxorubicin-related cardiotoxicity and vincristine-induced peripheral neuropathy.^37,38^ In our study, topotecan, navitoclax, venetoclax and idasanutlin were significantly more effective in tumor than non-malignant samples. Consistent with these findings, clinical studies report tolerability with promising clinical outcomes for topotecan, navitoclax and venetoclax in neuroblastoma or other pediatric solid tumors.^39–43^ Interestingly, topotecan also demonstrated high efficacy in both *ex vivo* and *in vivo* models. Although idasanutlin showed more efficacy in tumor than non-malignant samples, a phase I trial of idasanutlin in patients with leukemias or solid tumors was terminated due to toxicity and lack of efficacy.^44^ Non-malignant samples were more sensitive to mTOR inhibitors temsirolimus and everolimus. A clinical trial combining temsirolimus with re-induction chemotherapy reported excessive toxicity in children with relapsed acute lymphoblastic leukemia.^45^ However, interpretation is limited by the small sample size for non-malignant samples. In addition, this analysis considered all tumor samples collectively, irrespective of genetic or transcriptomic features, and all non-malignant samples irrespective of tissue origin. Future studies should focus on establishing larger reference panels including pediatric liver, kidney, and intestine tissues, as they are the key organs for drug metabolism and most affected by treatment toxicity.^46–49^ Including non-malignant pediatric tissues in *ex vivo* drug screening may help identify drugs that could cause unacceptable toxicity. This is particularly relevant in neuroblastoma, where young survivors face long-term treatment-related risks of (cardiovascular) complications and secondary malignancies.^50^

In conclusion, our findings demonstrate that drug sensitivities were generally consistent between short-term and organoid screens, but these models differ markedly in feasibility and turnaround times. Future studies will be essential to correlate *ex vivo* response with in-patient response. While organoids and PDX models remain essential for large-scale drug discovery and biomarkers research, *ex vivo* short-term screening offers high success rates and rapid establishment times suitable for application in functional precision oncology in neuroblastoma and possibly other pediatric and adult tumors.

## Methods

### PDX establishment at the ITCC-P4 consortium and KiTZ

Informed consent to participate in research and for the collection and use of biological samples was obtained from all patients, or their parents or legal guardians, in accordance with the Declaration of Helsinki.

The establishment and characterization of mouse Patient-Derived Xenografts (PDXs) used for drug screening in the Princess Máxima Center were performed by the ITCC-P4 consortium, as described before,^15–17^ and samples were expanded by EPO^51,52^ in Berlin, Germany, or Charles River in Freiburg, Germany. Before sample collection, studies were approved by the Ethics Board of the Medical Faculty of the University of Heidelberg or by the respective local authorities. Informed consent included permission for anonymized data sharing in academic and nonprofit settings, according to the Helsinki guidelines. Fresh tumor material (obtained through surgical resection or biopsy), a constitutional (germline) sample, and clinical information were collected for each patient. For the generation of PDX models, tumor tissue was subcutaneously transplanted to either NSG (NOD.Cg-Prkdc^scid^ IL2rg^tm1Wjl^/SzJ; Charles River Laboratories, Germany), Swiss Nude (Crl:NU(Ico)-Foxn1nu), CB17 SCID (CB17/Icr-Prkdc^scid^/IcrIcoCrl), or NOG (NOD.Cg-*Prkdc^scid^ Il2rg^tm1Sug^*/JicTac; Taconic Denmark) immunodeficient mice. All sample and clinical information, as well as analyzed molecular data of patient tumors, PDXs, and organoids) were barcoded and subsequently operated via R2 genomics analysis and visualization platform (r2-itccp4.amc.nl).

At KiTZ, patient material was collected and dissociated as described before, in agreement with local institutional ethical regulations, approved by the institutional review board, and with patient consent under de guidelines of the MAST protocol.^14^ Up to 1 million dissociated cells in PBS were mixed with Matrigel (1:1) and subcutaneously injected into the right flank of NSG mice under isoflurane anesthesia. Tumors were passaged three times in mice, and drug screening as well as PDX long term cultures were established from human-to-mouse passage 0 and/or mouse-to-mouse passage 3. All animal experiments were conducted in accordance with the German Animal Welfare Act and approved by local authorities.

### ITCC-P4 drug screen in PDX for response characterization

ITCC-P4 PDX models were characterized for their response to single compound or combination therapies in preclinical single-mouse trials.^16,17^ Based on individual formulation and specific schedules, single compounds and combinations were administered either orally, intravenously, subcutaneously, or intraperitoneally (Table 1). When the PDX tumor volume achieves a size of 100-250 mm³, a blinded randomization procedure was used to distribute mice to test groups with the subsequent start of treatment. The actual amount of compound was calculated daily depending on the individual body weight of animals throughout a four-cycle 28-day treatment phase. All animals were terminated at individual time points when they reached the maximum ethical tumor volume threshold (TV > 1.5cm³), body weight loss (>20%) limits, or when animals suffered from adverse effects before the end of four cycles. Control animals receive the most demanding schedule and therefore the lorlatinib vehicle formulation. From the first treatment day onwards, tumor size is measured twice a week until the end of the treatment.

**Table 1.** The treatment schedule of the compounds used in vivo, with the compound name, the dose given to the mouse, the schedule, the route of administration, and the formulation of the given compound. On represents days on treatment, off represents days off treatment.

| Drug | Dose mouse<br>(mg/kg) | Schedule | Route | Formulation |
| --- | --- | --- | --- | --- |
| **Etoposide** | 5 | Daily 5-on/2-off for 4<br>cycles | Intravenous | 0.9% NaCl |
| **Topotecan** | 0.5 | Daily 5-on/2-off for 4<br>cycles | Intravenous | 5% Glucose |
| **Idasanutlin** | 100 | Twice a day,<br>5-on/2-off for 4<br>cycles | Orally | GTX011795 (water 97.8%,<br>2% hydroxypropyl<br>cellulose, 0.1%<br>polysorbate, 0.09% methyl<br>paraben, 0.01% propyl<br>paraben, 0.0072% sodium<br>acetate, 0.0568% acetic<br>acid |
| **JQ-1** | 50 | Daily, 5-on/2-off for<br>4 cycles | Intraperitoneally | Vehicle: 1:10 DMSO: 10%<br>hydroxypropyl-β-<br>cyclodextrin |
| **Prexasertib** | 10 | Twice a day, 3-on/4-off for 3 cycles | Subcutaneously | 20% captisol in purified<br>water |
| **Cobimetinib** | 7.5 | Daily, for 28 days | Orally | 0.5% methylcellulose/<br>0.2% Tween 80 (MCT) |
| **Lorlatinib** | 3 | Twice a day, for 28<br>days | Orally | MilliQ DI water, 1:1.2<br>lorlatinib:HCl molar ratio |
| **Copanlisib** | 10 | Daily, 2-on/5-off<br>4 cycles | Intravenously | 0.9% NaCl |
| **Alectinib** | 20 | Daily, for 28 days | Orally | 5% DMSO, 5% cremophor<br>EL solution, 90% NaCl<br>(0.9%) |
| **Belvarafenib** | 15 | Daily, for 28 days | Orally | 10% DMSO, 10%<br>cremophor, 15% PEG400,<br>15% HPCD, 0.02N HCl in<br>sterile water |

### Tissue dissociation

PDX-derived tumor tissue (both cohorts) and patient-derived tumor tissue (KiTZ) were processed using a site-specific enzymatic digestion protocol. Tissue was incubated in Liberase DL (Sigma Aldrich, cat. no. 5466202001; 1:43 dilution) in tumor stem media (TSM)^14^ at 37°C for 30 minutes with occasional swirling (Máxima) or in trypsin (10 mg/ml) at 37°C for ten minutes (KiTZ). TSM consists of 47.5% Neurobasal-A medium, 47.5% D-MEM/F-12, 1% HEPES buffer solution (11M), 1% sodium pyruvate MEM (1001mM, CE), 1% MEM non-essential amino acids solution (101mM), L-glutamine solution BIOXTRA (21mM), and 1% penicillin-streptomycin-glutamine (all from Life Technologies, Thermo Fisher Scientific Inc.). This was supplemented with 2% B27 Supplement Minus Vitamin A (50×; Life Technologies, Thermo Fisher Scientific Inc., Waltham, MA, USA), 0.02% (w/v) recombinant human EGF (100 µg/mL), 0.02% (w/v) recombinant human FGF-basic (100 µg/mL), 0.05% H-PDGF-AA (20 µg/mL) (all from PeproTech Inc., Rocky Hill, NJ, USA), and 0.1% heparin solution (21mg/mL; Sigma-Aldrich, Munich, Germany). Enzymatic activity was stopped by harvesting the supernatant, adding medium (Máxima), or by adding 12 µl trypsin inhibitor, followed by repeated additions of a 1:1 mixture of 1 M MgCl_2_ and DNase (2 mg/mL; 60 µL per addition) until the tumor fragment settles (KiTZ). At the Máxima, the residual tissue underwent a second digestion in Liberase TM (Sigma Aldrich, cat. no. 5401119001; 1:43 dilution in TSM). After another 30 minutes, cells were harvested as described above. Cells were filtered through a pre-washed 70 µm cell strainer (Máxima) or a 70-or 100 µm strainer (KiTZ) and centrifuged at 300 g for 5 minutes (Máxima) or at 400-500 g for 10 minutes (KiTZ). Red blood cells were lysed using lysis buffer with a maximum incubation time of 2-5 minutes at room temperature. Cells were resuspended in TSM complete^14^ and counted using trypan blue.

### Short- and long-term organoid culture

After tumor tissue dissociation, cells were cultured for short-term or until organoids were established. In all cases, tumor cells were cultured at 37°C, 5% CO₂. Short-term cultures and screens at the Máxima and KiTZ were done in TSM. At the Princess Máxima Center, DNA was harvested before drug screening. Freshly dissociated cells were cultured for 5-8 days (at Máxima) or for 2-7 days (depending on growth and morphology, at KiTZ), before screening. Two short-term screens were excluded due to mice contamination.

Long-term organoid cultures at the Máxima were cultured in Neuroblastoma Organoid Medium (NBOM), consisting of DMEM with GlutaMAX™ (Life Technologies, cat. no. 21885108), 25% Ham’s F-12 Nutrient Mix, B-27™ Supplement (minus vitamin A), N-2 Supplement, 100 U/mL penicillin, 100 µg/mL streptomycin, 20 ng/mL recombinant human EGF, 40 ng/mL recombinant human FGF-basic, 200 ng/mL recombinant human IGF-I, 10 ng/mL recombinant human PDGF-AA, and 10 ng/mL recombinant human PDGF-BB.^53^ At KiTZ, long-term cultures were cultured in TSM. In both centers, cells were cultured biweekly, with medium refreshment or passaging based on microscopic evaluation. Medium was refreshed by harvesting medium, centrifuging at 250 g for 5 minutes, and resuspending the pellet in fresh medium. Oversized cultures with necrotic cores were dissociated by pipetting in a 50 mL tube before splitting (Máxima) or using Trypl Express (Life Technologies, Thermo Fisher Scientific Inc., Waltham, MA, USA; 5 min, 37 °C; KiTZ). Mycoplasma testing and STR profiling (PowerPlex system and GeneMapper software, Promega) were performed routinely. Drug screening of long-term organoid cultures was initiated only after stable growth of human tumor cells was confirmed. Tumor purity was determined by comparing patient-specific breakpoints of the culture (using SNP array) to those of the original tumor. On the day of organoid screening, cryovials, RNA, and DNA were harvested at the Máxima.

Part of the KiTZ short-term screens were previously reported in Peterziel *et al.*^14^, Máxima patient material organoids were previously reported in Langenberg *et al*.^9^, and Máxima short-term screens on *ex vivo* cultures from PDX models were previously reported.^15^

### Molecular profiling

For models used for drug screening in the Máxima, comprehensive molecular profiling was performed as part of the ITCC-P4 consortium^16,17^ in Paris (Curie) and Heidelberg (KiTZ), on PDX and organoid samples, and if available, on primary patient tumors and matched germline controls. For models used for drug screening at KiTZ, molecular profiling was exclusively performed on the primary patient tumor. For low-coverage whole-genome sequencing (lcWGS) and whole-exome sequencing (WES), libraries were prepared using standardized methods^54^ and captured with the Agilent SureSelect Clinical Research Exome V2 Kit, followed by sequencing using Illumina platforms at both centers. For PDX models, organoids and, if available, primary patient tumors, used for drug screening at Máxima, RNA-seq was also performed by the ITCC-P4 consortium^16,17^ in Paris (Curie) and Heidelberg (KiTZ). RNA-seq was performed on the NextSeq 500 (150 nt paired-end) or NovaSeq (100 nt paired-end).

### Drug libraries

The Máxima drug libraries L13 and L14 consist of 224 and 226 drugs, respectively, including FDA-approved agents and those in (pre-)clinical development (Suppl. Table 1). All organoids are screened with the full library. For short-term screens, the full library is used when sufficient biomass is available; otherwise, the clinical subset of 134 drugs is used. The KiTZ drug library consists of 78 clinically relevant anticancer drugs, including targeted as well as chemotherapy (Suppl. Table 2). Both libraries consist of clinically relevant anticancer drugs, including FDA-approved agents and those in (pre-) clinical development. Most drugs are dissolved in DMSO and stored under nitrogen at room temperature (Máxima) or in an oxygen- and moisture-free environment at room temperature (KiTZ). At the Máxima, exceptions include five drugs (metformin, perifosine, carboplatin, oxaliplatin, and copanlisib) dissolved in MQ, and cisplatin in saline, which are stored at –20 °C. Before screening, 384-well working plates (Labcyte 384LDV) are shaken (30 min at room temperature) and centrifuged (2.5 min at 1500 rpm). The Echo 550 dispenser is then used to verify well volume (>2.5 µL) and DMSO percentage (>80%).

### Drug screening

Drug screens were performed in duplicate. Cells (5,000/well in 40 µL at the Máxima; 1000/well in 25 µL at KiTZ) were plated into 384-well flat-bottom tissue culture plates (#3764, Corning) using a multidrop combi dispenser (Thermo Scientific) and cultured for 24 hours (Máxima) or directly screened (KiTZ, as described in Peterziel *et al.*^14^). At the Máxima, cell viability was assessed at t = 0 using the Cell Titer Glo 3D (CTG3D) assay (Promega, G9683) according to the manufacturer’s protocol.

Using the Echo 550, 100 nL of each drug (in DMSO or MQ) was added to achieve final concentrations ranging from 0.1 nM to 100 µM (0.25% DMSO/MQ). DMSO-only treated cells served as positive controls; 10 µM staurosporine-treated cells were used as negative controls. Cells were incubated with the drugs for 72 hours at 37°C and 5% CO₂, except for organoid screens at the Máxima (120 hours).

Data were normalized against DMSO-treated (100% viability) and empty well controls (0% viability). Dose-response curves were modeled using a five-parameter log-logistic model with the R package drc (v3.0.1; R 4.3.1^55^). From these models, the IC50 (concentration for 50% viability reduction), area under the curve (AUC), and relative cell growth (Máxima, CTG3D signal day 5/t0) were calculated. Screen quality was assessed based on growth dynamics, controls, and consistency across replicates. AUC was the primary metric used for defining and comparing drug sensitivities, as IC50 could not be reliably determined for inactive drugs. We normalized the area under the curve (AUC) between 0-100, accounting for differences in the number of concentrations across cohorts.

### Calling of genomic alterations

Whole-exome sequencing was used to identify genomic alterations as described above. High-level amplifications of *MYCN*, *ALK*, *CDK4*, *KRAS*, and *MDM2* were called when copy number (CN) > 8. ALK alterations were also called in case of an activating mutation within the tyrosine kinase domain. Alterations in genes with suppressor gene function *TERT*, *NF1*, *PTPN11*, *PIK3CA*, CDKN2A/B and *AKT* were considered based on homozygous deletions, or loss of heterozygosity (LOH) in combination with an inactivating mutation. ATRX was called in case of homozygous inactivation or deletion, or the combination of LOH, in-frame fusions, inactivating mutations, or hotspot mutations. TP53 mutations were called in case of a deletion combined with an inactivating (frameshift or SNV) mutation. Chromosomal gains were called when CN > 2.5 and losses when CN < 1.5.

### RNA-sequencing normalization and gene set enrichment

RNA-sequencing data were aligned to the human reference genome (GRCh37, GENCODE 19) using STAR^56^ and the raw counts were normalized to CPM values using the Trimmed Mean of M-values (TMM) method implemented in the edgeR^57^ R package (version 4.6.2) and log2 transformed. Gene set scores were calculated using GSVA^58^ (version 2.2.0) and gene sets were obtained from the Reactome database.^59^

### Statistical testing

Statistical analysis was performed using the Wilcoxon rank-sum test.^60^ Correlation was calculated using Pearson correlation^61^ (package stats, version 3.6.2) to assess linear relationships. For in vivo-ex vivo comparison, Spearman concordance was used. P-values were used to evaluate the statistical significance of the observed correlations. Differential response analysis between tissue sources and models was performed using Limma^62^ (version 3.21). To correct for multiple hypothesis testing, p-values were adjusted using the Benjamini-Hochberg^63^ method. Significance was considered when the p-value was below 0.05.

## Acknowledgements

We thank all patients and their parents for providing samples that made this research possible. We appreciate the support given by the High Throughput Screening Facility in the Princess Máxima Center and our collaborators of the ITCCP4 consortium for sharing their knowledge and resources. The work performed in the Maxima was supported by the IMI2 ITCC-P4 program (grant agreement no. 116064). Furthermore, we would like to thank Marko Weidmann, Anette Hugo, Alexandra Stroh-Dege, Ina Siebig and Carina Müller for their excellent technical assistance. We thank Laura Turunen and Jani Saarela from the High Throughput Biomedicine Unit and Vilja Pietiäinen (Institute for Molecular Medicine Finland, Helsinki Institute of Life Science, University of Helsinki, Helsinki, Finland and Biocenter Finland) for the support with drug-plate printing and the provision of ready-to-go drug plates. We thank the INFORM program for supporting us with patient-derived fresh tissue.

## Data availability statement

The datasets used and/or analysed during the current study available from the corresponding author on reasonable request.

## Competing interest statement

IO received research grants from PreComb, BVD and Day One Therapeutics. GS receives research funding from Roche, BMS, Pfizer and MSD. The other authors declare no competing interests related to this study.

## Author contributions

Marlinde. C. Schoonbeek: conceptualization, data curation, analyses, investigation, methodology, writing-origional draft, visualisation, project administration. Marloes van Luik: data curation, analyses, writing-origional draft, visualisation. Heike Peterziel and Dennis Gürgen: investigation, conceptualization, writing – review & editing. Vicky Amo-Addae: methodology, recources (Máxima library). Lindy Vernooij: project administration. Eleonora J. Looze and Sabina Valova: investigation. Sina Kreth, Angelika Eggert, Julia Schueler, Frank Westermann: methodology, recources (PDX models). Jan Koster: resources (R2 platform, molecular profiling). Aniello Federico, Apurva Gopisetty, Benjamin Schwalm, Marcel Kool: methodology, data curation, analyses, recources (molecular profiling). Alexandra Saint-Charles: project administration. Gudrun Schleiermacher: methodology, recources (molecular profiling and PDX models). Gilles Vassal, Hubert N. Caron, Lou Stancato, Stefan M. Pfister: conceptualization, supervision (ITCC-P4 leadership). Jens Hoffman: investigation, supervision. Arjan Boltjes: visualization, supervision. Ina Oehme: conceptualization, supervision, writing – review & editing. Marlinde L. van den Boogaard, Selma Eising, Jan J. Molenaar, Sander R. van Hooff: conceptualization, supervision, writing – review & editing. All authors reviewed the manuscript.

