## Supplemental Figures and Information for "Sensitivity profiling reveals consistent drug responses across preclinical neuroblastoma models"

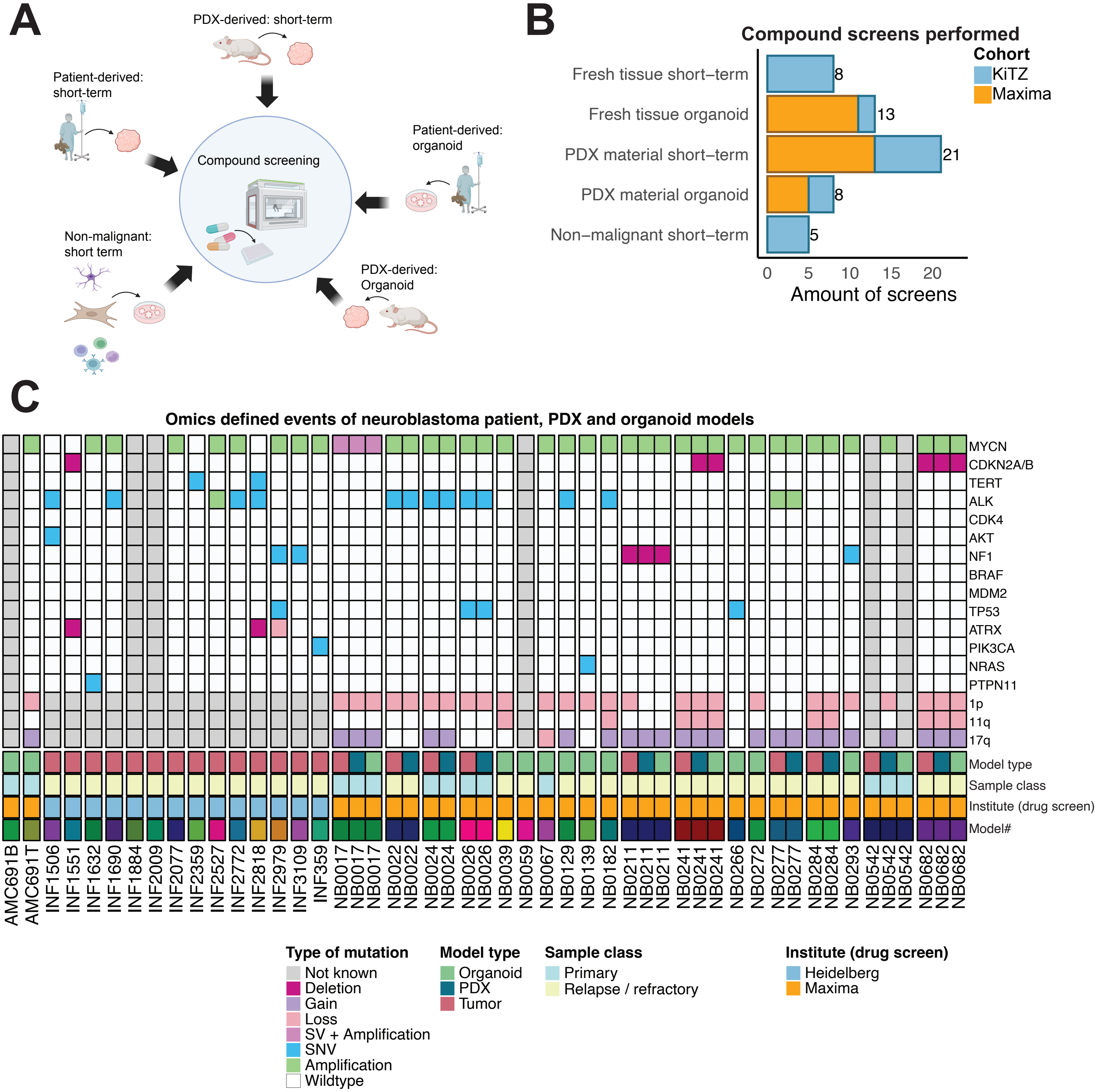


***Supplementary Figure 1. Establishment and genetic characterization of models for compound screening. A)*** *Five model types were generated across centers: patient-derived short-term and organoid models, PDX-derived short-term and organoid models, and short-term screens of non-malignant samples.* ***B)*** *Bar plot representing the total number of compound screens per model type of the Máxima Center and KiTZ. Part of the short-term and organoid screens performed at KiTZ were previously described in Peterziel et al., Máxima patient material organoids were previously described in Langenberg et al., and short-term screens performed at Máxima in Schoonbeek et al.* ***C)*** *Oncoplot of genomic landscape of neuroblastoma samples, showing key gene alterations and chromosomal aberrations colored by alteration type. Institute (drug screen) refers to the institute where the drug screen is performed. PDX = Patient-Derived Xenograft, SNV = single-nucleotide variant, SV = structural variant.*

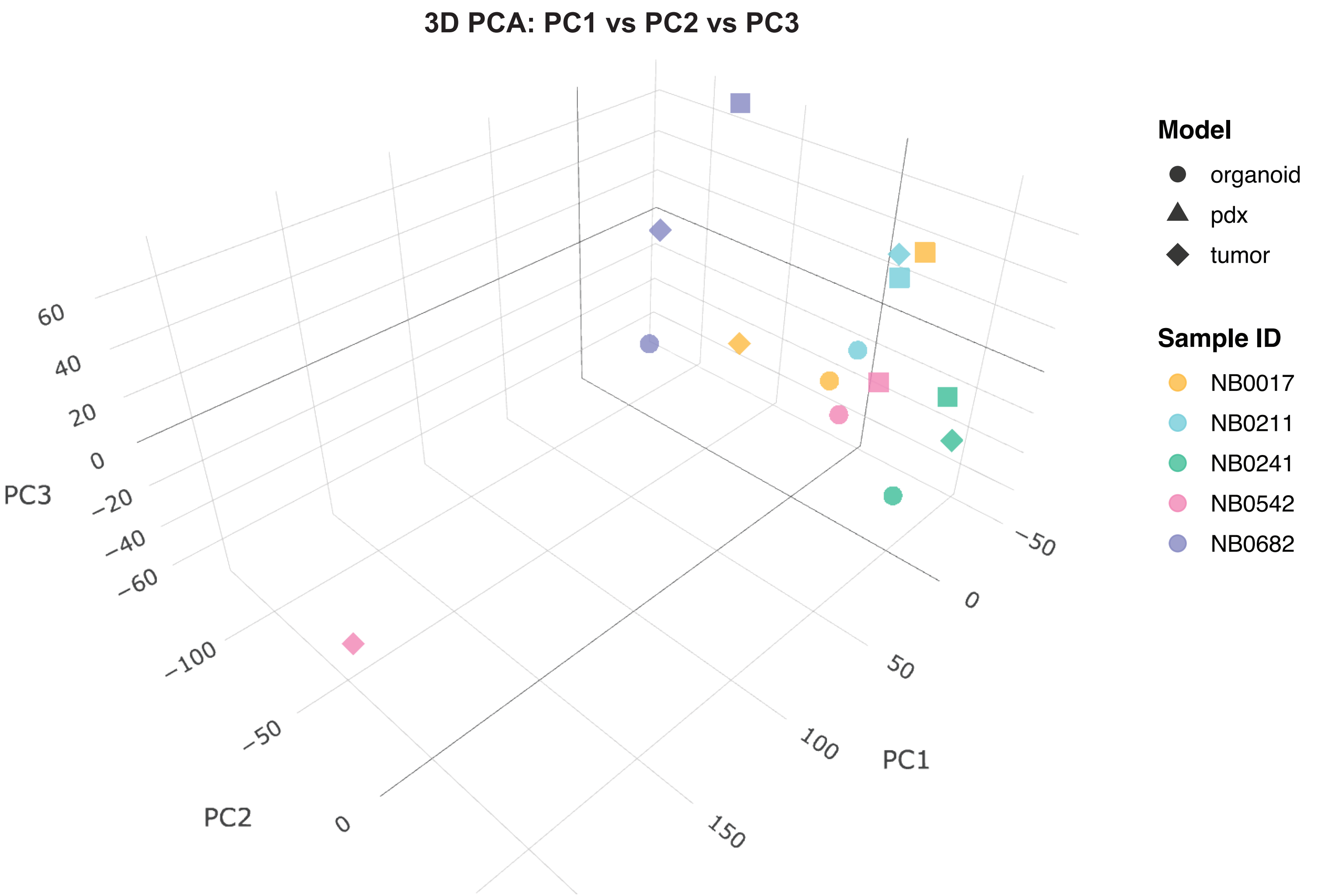


***Supplementary Figure 2. Transcriptomic profiling across matched organoid, PDX and patient samples.*** *Three-dimentional principal component analysis (PCA) plot of gene expression profiles from matched neuroblastoma patient tumors, patient derived xenografts (PDX) and organoid models. Principal components 1, 2 and 3 (PC1-3) are shown. Each point represents an individual sample; shapes indicate the model (type organoid, PDX or patient) and the colors indicate matched sample IDs.*

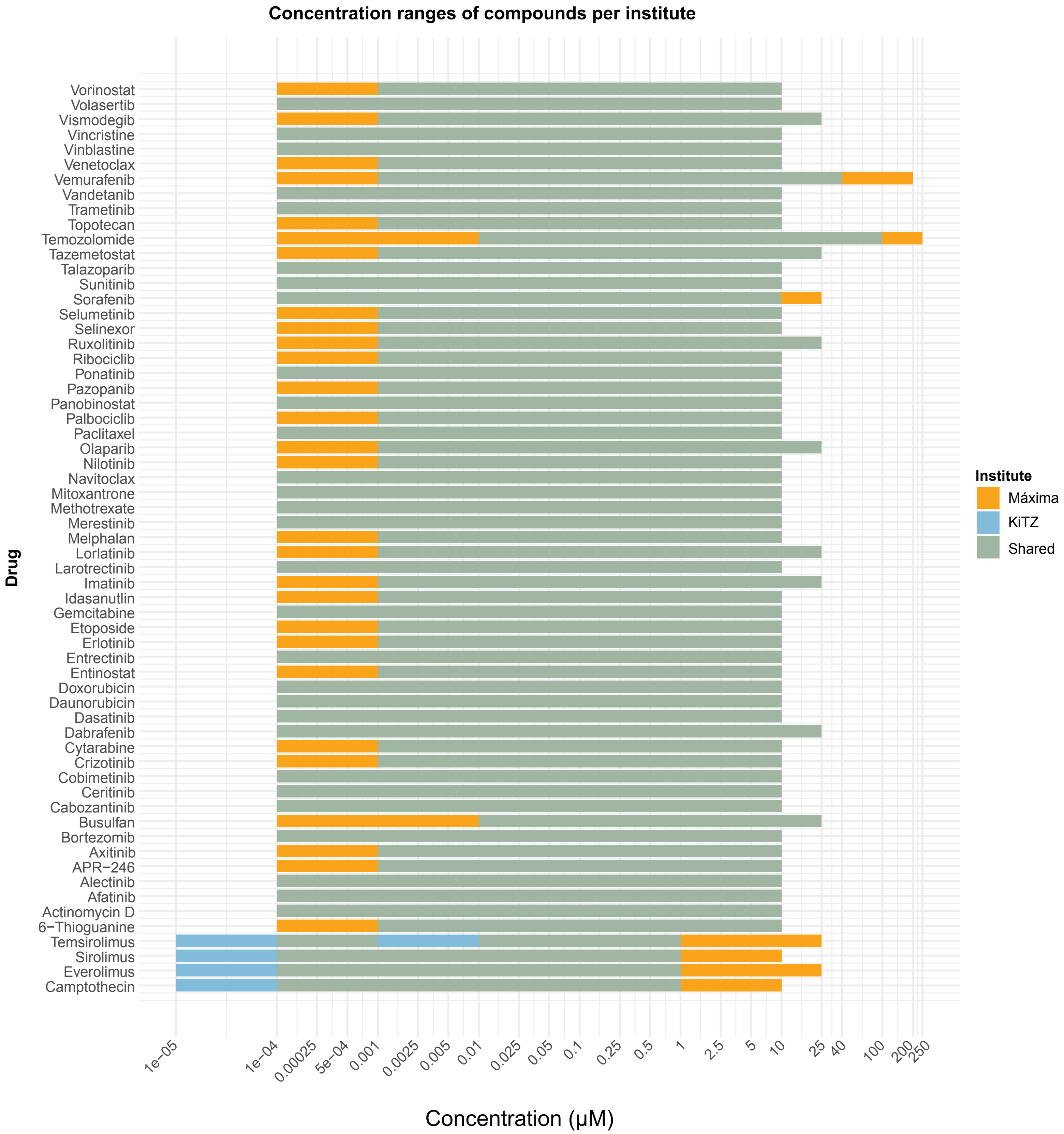


***Supplementary Figure 3. Concentration ranges of the overlapping drugs per institute.*** *Concentrations of drugs in Máxima (orange), KiTZ (blue), and overlapping (green). Concentrations are between 1 x 10-6 μM to 250 μM.*

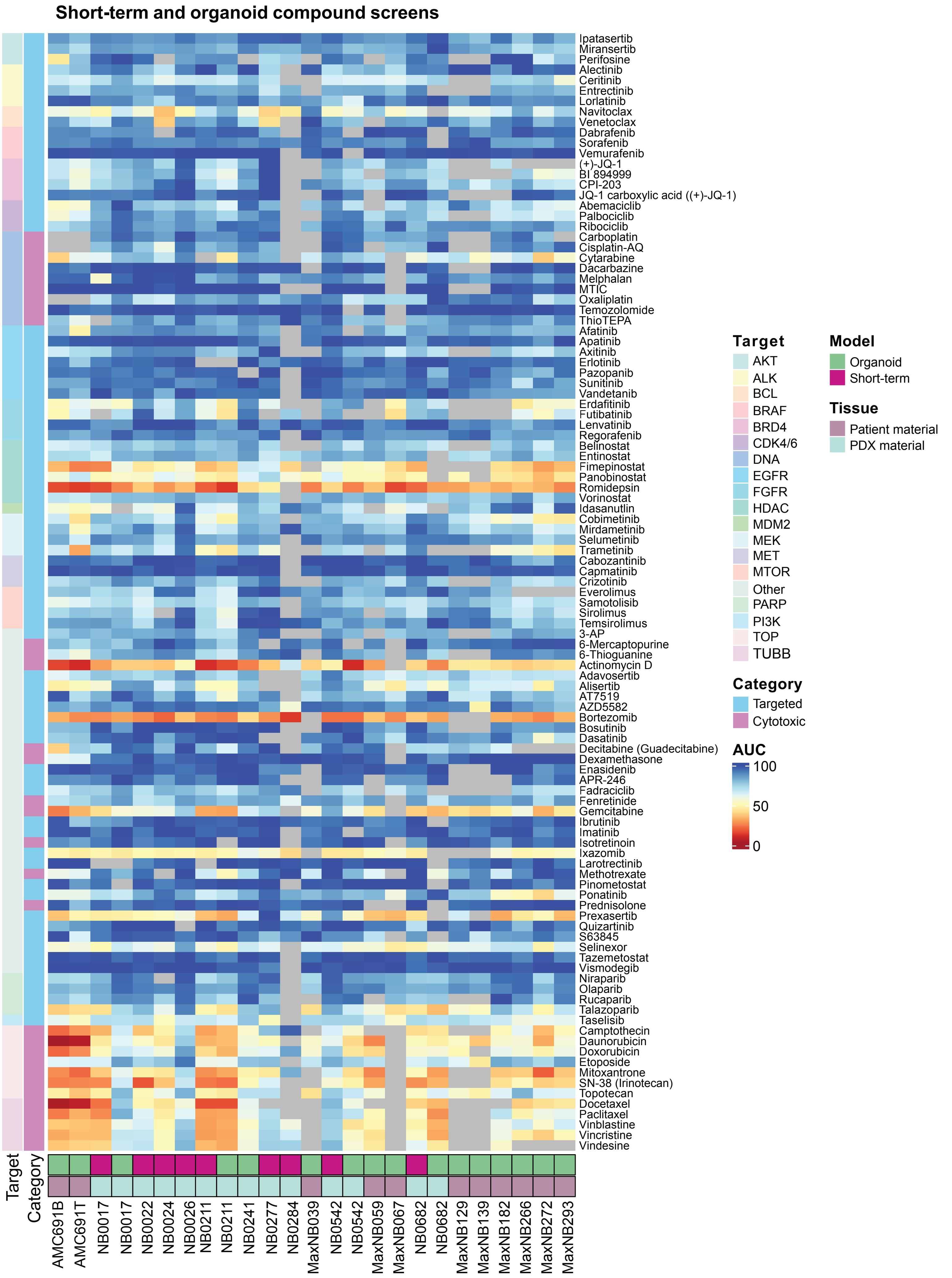


***Supplementary Figure 4. Compound screening on Máxima neuroblastoma samples.*** *The normalized area under the curve (AUC) of the short-term and organoids screens of the Máxima cohort, lower AUC values indicates more sensitivity to the compound. Samples are colored on screen type and tissue origin of the sample. Compounds are colored based on target and category of the compound (cytotoxic or targeted). Compounds of the clinical priority library are shown as they are screened in most samples.*

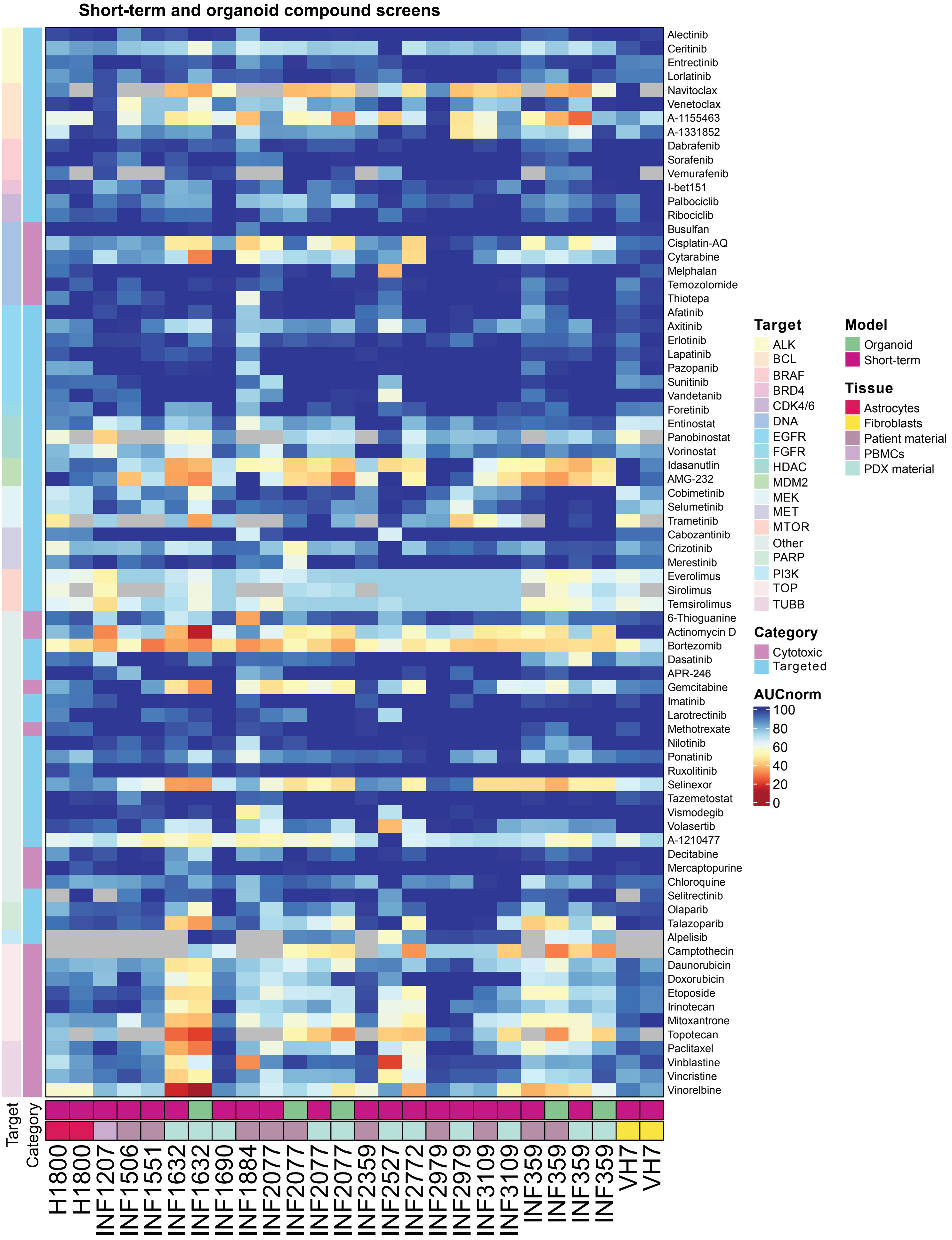


***Supplementary Figure 5. Compound screening on KiTZ neuroblastoma and non-malignant samples.*** *The normalized area under the curve (AUC) of the short-term and organoids screens of the KiTZ cohort. Samples are colored on screen type and tissue origin of the sample. Compounds are colored based on target and category of the compound (cytotoxic or targeted).*

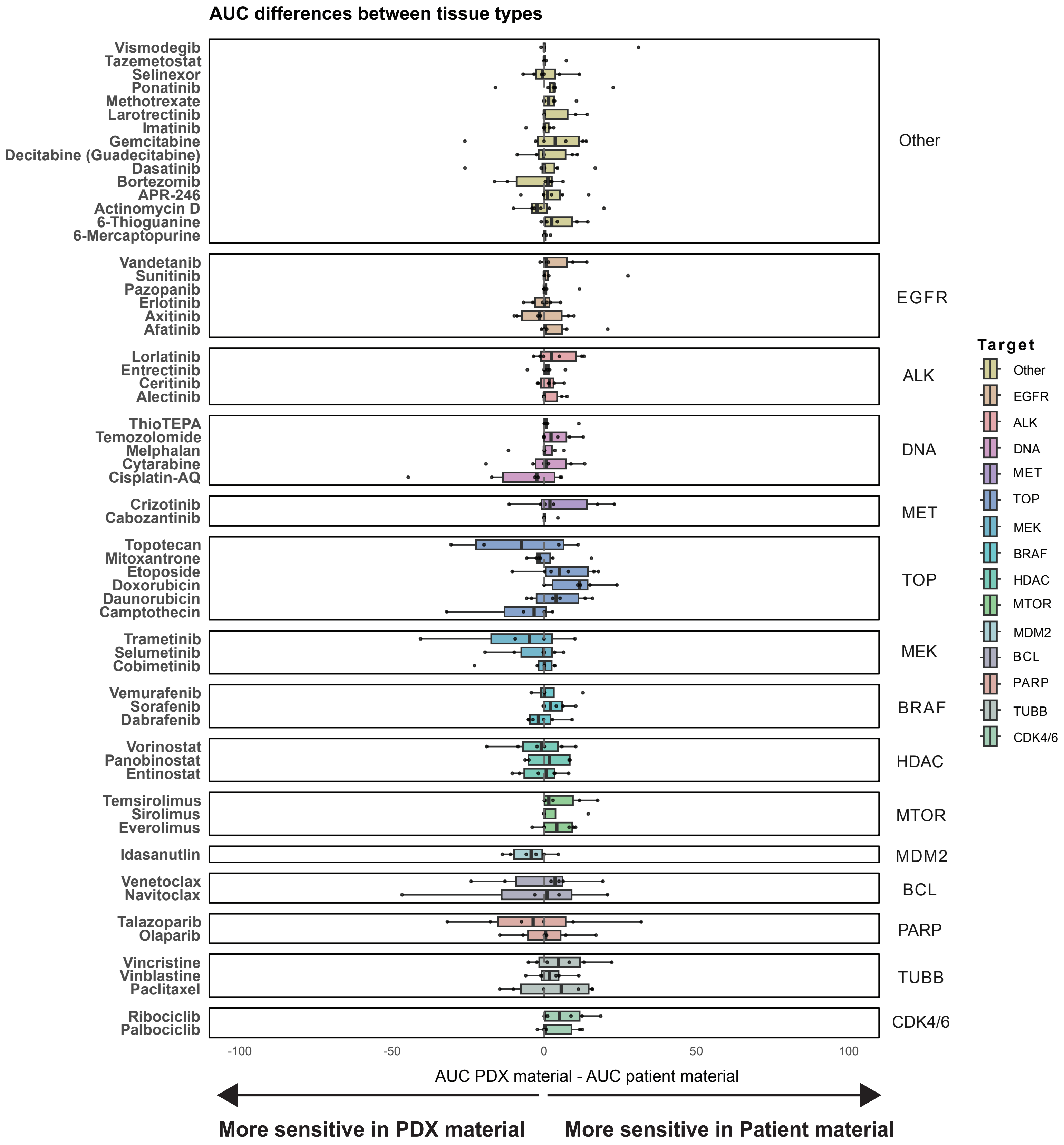


***Supplementary Figure 6. Differences in drug sensitivity between screens from patient material and PDX material of matched samples.*** *For each sample pair, the AUC of the PDX material screen is substracted by the AUC of the patient material screen. Positive numbers indicate higher sensitivity in the patient material screen, negative numbers indicate higher sensitivity in the PDX material screen. Boxes represents the distribution over the differences per compound, colored per target. Each dot represents the difference between a sample pair.*

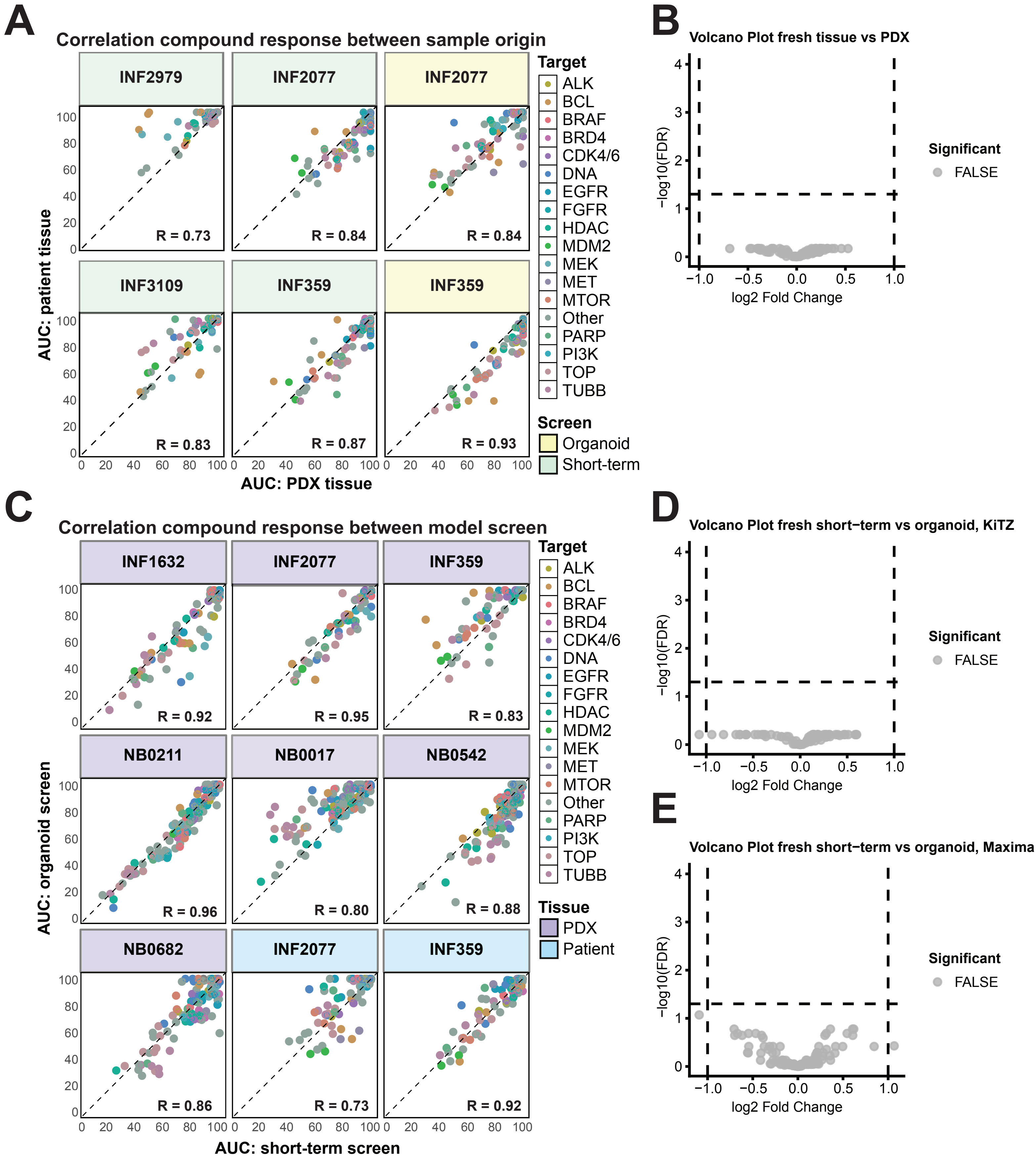


***Supplementary Figure 7. Comparison of drug response across neuroblastoma model systems and sample origin. A)*** *Correlation plots of compound response (AUC) on organoid or short-term screens from matched samples derived from different tissue origins (patient or PDX material).* ***B, D-E)*** *Volcano plot showing the influence of tissue origin (B: patient versus PDX) or screen type (D-E) on compound efficacy. A significant (adjusted p < 0.05; indicated by the dashed line) compound would mean that the AUC of that compound is influenced by tissue or screen type. In all cases, no compounds are found to be significant. Differential response analysis between tissue sources and models was performed using Limma (version 3.21).* ***C)*** *Correlation plots of the compound response (AUC) of organoids and short-term screens established from matched samples. In the correlation plots, each dot represents a compound, colored by its target. A lower AUC defines higher sensitivity. R represents Pearson correlation.*

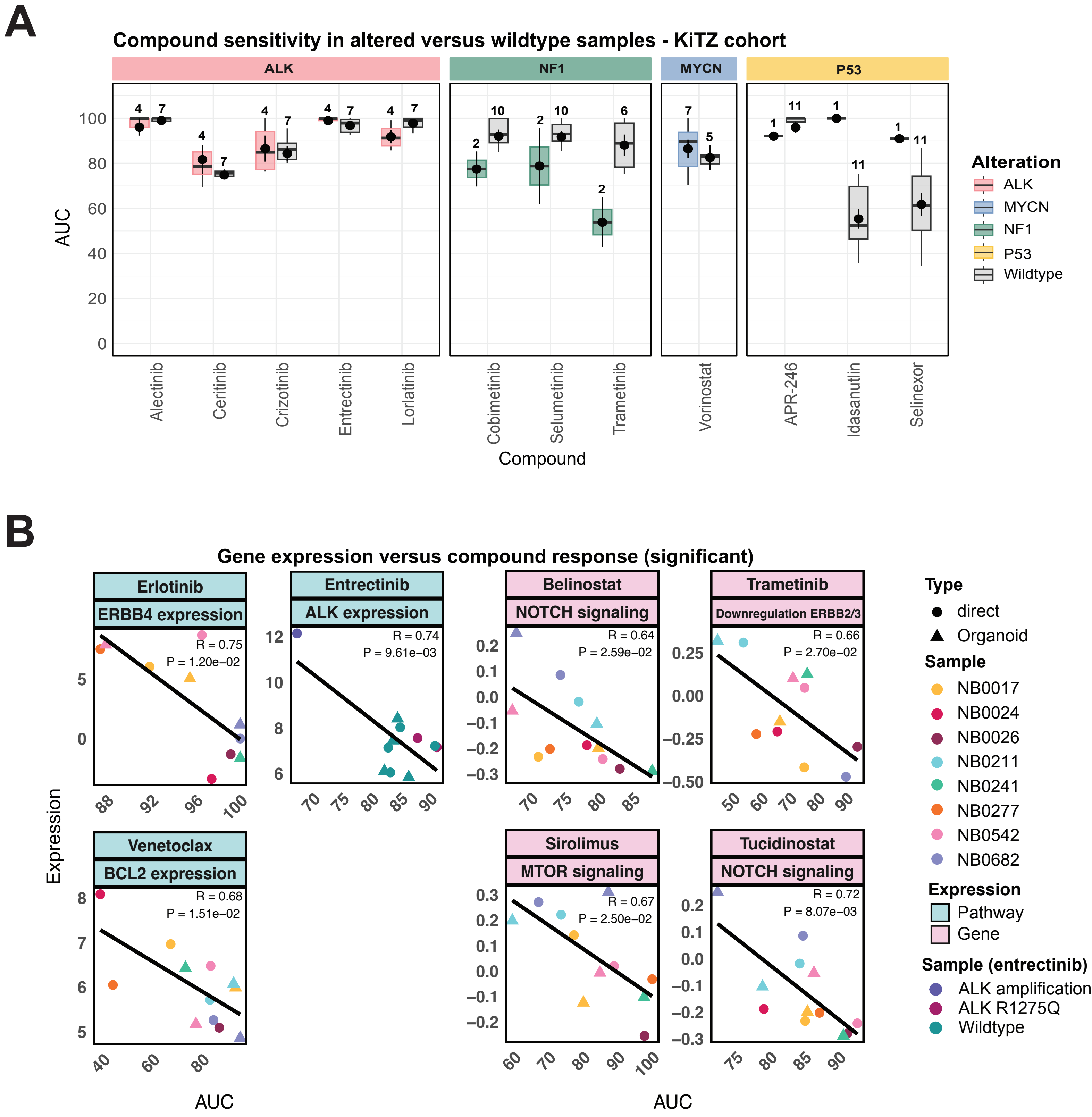


***Supplementary Figure 8. The association of genomics and transcriptomics with compound response. A)*** *Area under the curve (AUC) values of samples with targetable gene alterations, in boxes colored by alteration, compared to wildtype samples, in grey boxes, of the KiTZ cohort. The NF1/RAS altered samples consist of two NF1 altered samples (KiTZ). A lower AUC defines higher sensitivity to the compound. The dot indicates the mean, and the vertical line indicates the standard error around the mean. Stars indicate a significant difference between the altered and wildtype samples (adjusted p-value < 0.05).* ***B)*** *Correlation between AUC and expression of target/targeted pathway of compounds. Each dot represents a sample, colored by its name and shaped by model type. An exception is entrectinib, hwere samples are colored based on ALK-status. R represents the Pearson correlation. All correlations are significant (adjusted p-value < 0.05).*

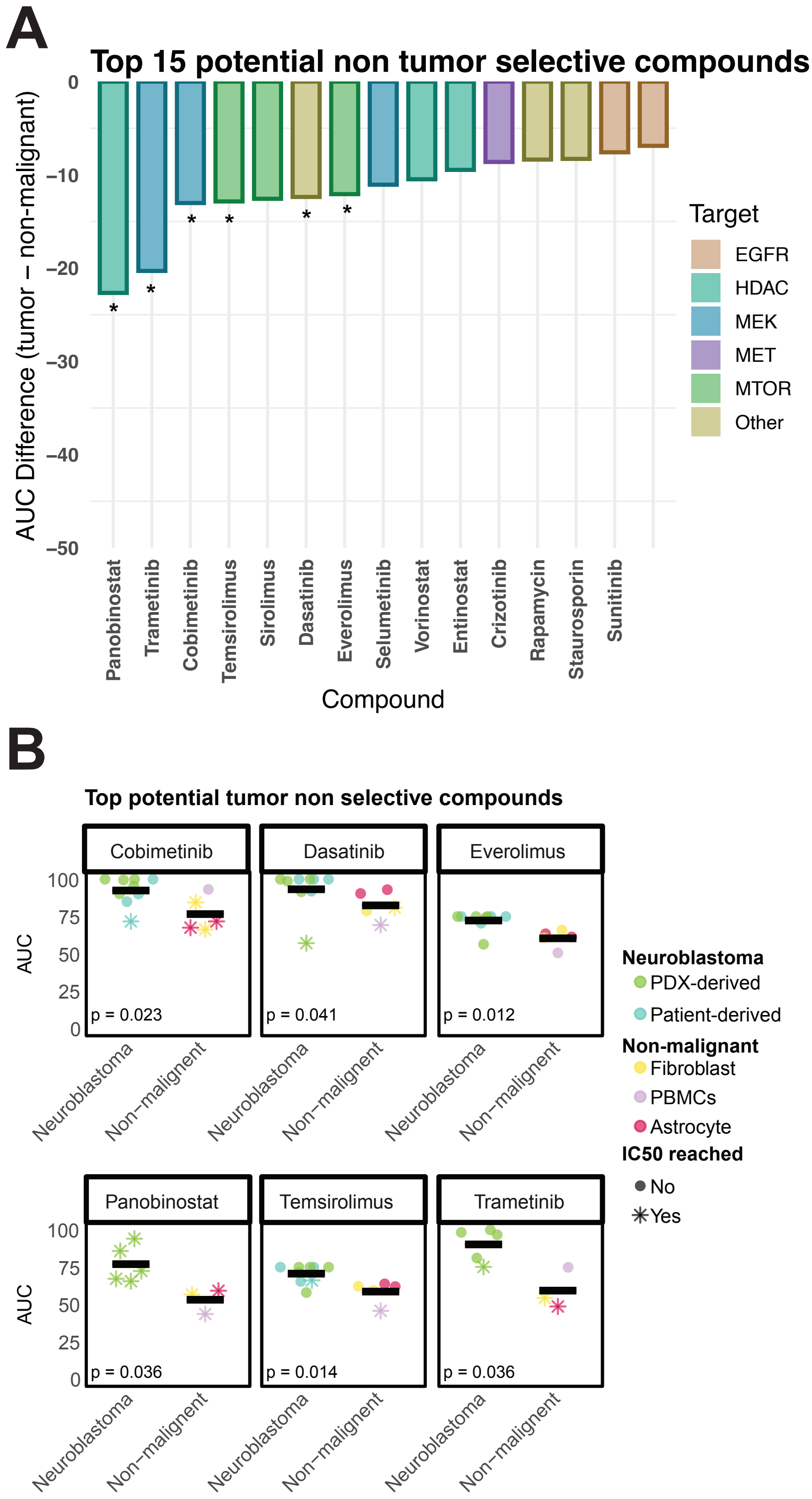


***Supplementary Figure 9. Potential (non) tumor selective compounds.***

***A)*** *Bar plot representing the top 15 tumor non-selective compounds. The top 15 compounds with the highest mean AUC difference between non-malignant and tumor samples (mean AUC non-malignant – mean AUC tumor) are shown. Bars are colored based on the target of the compound. Asterisks indicate that the compound is significantly more effective in non-malignant samples.*

***B)*** *Box plots representing drug sensitivity (AUC) for tumor and non-malignant samples for compounds that were more effective in non-malignant cells than in neuroblastoma. Displayed are compounds for which tumor samples had a significantly higher AUC than non-malignant samples and that ranked among the top 15 compounds with the largest mean differences between the two groups. Each dot represents the response of one sample, colored by tissue type. The horizontal line indicates the mean AUC of the samples.*

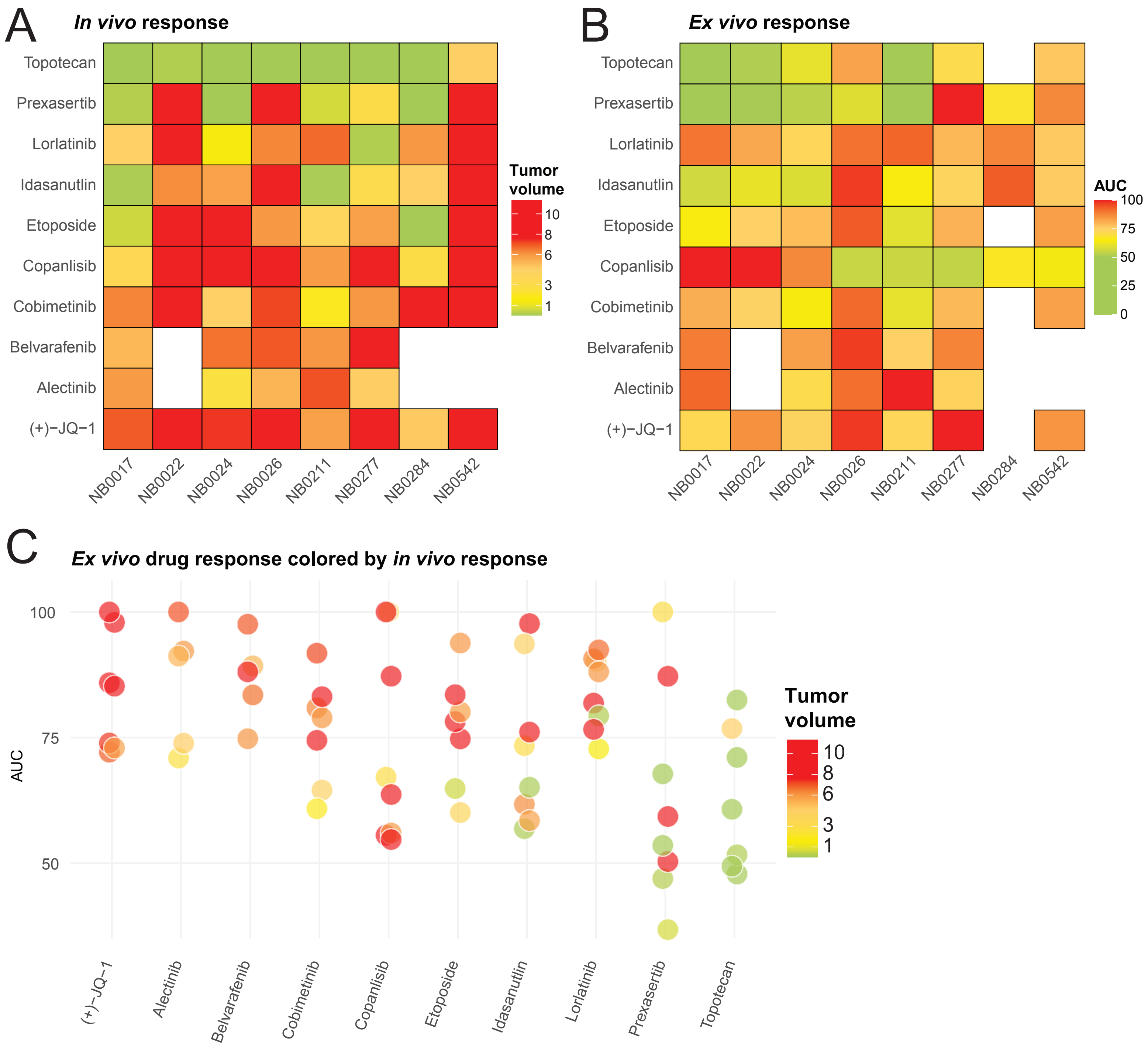


***Supplementary Figure 10. Comparing ex vivo and in vivo drug responses. A)*** *Dot plot that represents ex vivo response (area under the dose-response curve, AUC), colored by in vivo end-of-treatment response (Tumor volume, normalized to starting volume). Each dot represents a matched sample that is successfully tested both in vivo and in vitro. Higher sensitivity to the compound is defined by a lower AUC or a lower Tumor volume.* ***B)*** *Heatmap representing the in vivo treatment responses in PDX models. Each block represents a model (ordered by PDX model), colored by end-of-treatment response (Tumor volume, normalized to starting volume).* ***C)*** *Heatmap representing the ex vivo treatment responses of short-term cultured PDX models. Each block represents a model (ordered by sample), colored by AUC.*

### Supplementary information

**Supplementary Table 1**
Table representing the drug library that was used for screens at Máxima, including info on drug category, drug targets and pathways.

**Supplementary Table 2**
Table representing the drug library that was used for screens at KiTZ, including info on drug targets.

**Supplementary Table 3**
Table representing the overlapping drugs of the drug libraries in Máxima and KiTZ, including info on drug targets.

**Supplementary Table 4**
Table representing compounds that were tested in *in vivo* PDX models and the specific models per compound.
